# Mating pattern, duration and multiple mating in *Chondromorpha severini* Silvestri (Diplopoda: Polydesmida)

**DOI:** 10.1101/2020.08.23.263863

**Authors:** S. Bhakat

## Abstract

Mating behaviour of *Chondromorpha severini*, a polydesmid millipede was studied in the field and in the laboratory condition. Copulating pair follows the general rule of love play before actual act of coitus. Mating duration varied from one to 25 minute with an average of eight minute. Mating frequency was maximum in early and late hours of day. In the multiple mate preference experiment, 10 pairs of male and female were used to calculate preference index (P_i_) of individual sex. Preference index varies from 0.65 to 0.91. The implication of multiple mating has been discussed in detail. The study confirmed that i) the species belongs to polygynandrous mating system where males are the pursuers and females are the accomplishers ii) short and long duration mating is related to mate acquisition and mate guarding respectively

## Introduction

*Chondromorpha severini*, Silvestri is a common millipede in leaf litter of West Bengal (Mukherjee, 1962; Bhakat, 2014).The millipedes are abundant in the months of June to September when the males and females are pairing. An attempt has therefore been made in the present investigation to study the different aspects of sexual behaviour like pattern of mating, seasonal variation of mating, mating duration, hour of mating etc. of the species in the natural surroundings as well as in the laboratory.

## Materials and methods

Field studies were made at an undisturbed site (50 m × 12m) surrounded by trees and containing rotten wood, bark and leaf litter in Suri (87°32’00’’E, 23°55’00’’N). Initially for 10 days (3.VII.2018 to 13.VII.2018) the number of mating pairs at every hour of the day and night was noted. Since no mating during the night was observed, subsequent observations were confined to day time. During each month from May 2017 to April 2018, observations were taken at weekly intervals. On each observation, the numbers of mating pair were recorded.

Observation on mating frequency, love play and mating were made in the laboratory, keeping adult millipedes in glass troughs (dia. 30cm, height 12cm) filled with soil and decomposed litter collected from their natural habitats. There were 10 such glass troughs, each containing seven pairs (seven males and seven females) of adults. Mating duration was measured from the moment the pair assumed the characteristics mating position (ventro-ventro) till the time when the male and female are separated.

Preferential mating was studied with 10 pairs (10 males and 10 females) of adults whose activities were followed for five consecutive days. Millipedes were individually marked on the dorsum with plastic paints. The pairs were kept under scrutiny from 0600 to 1830 hours. Each pair was housed in a small plastic jar (dia. 10cm, height 10cm) filled with soil and decomposed leaf litter. There were 10 such troughs marked as A, B, C, …..I, J. After three hours of observations (either in fore- or afternoon) partner of each was changed e. g. by transferring male from A to B, B to C and so on. So each of 10 male had opportunity to meet with each of 10 females. To evaluate mate preference of male or female, an index is formulated named as Preference Index (P_i_) (of a particular sex) which can be calculated by:

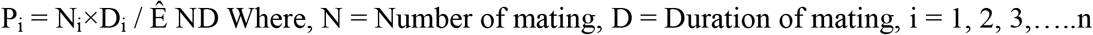

## Results

### Love play

*C. severini* was found to practice love play before the actual act of mating. During this love play the male approached the female from posterior and gradually mounted on her back and proceeded anterior and came to her head region. He then vigorously moves his legs (just like dance) for a few seconds. Thereafter he rubs his legs on the dorsal and lateral sides of the female. The receptive female then engaged herself in mating within a minute or two.

If the female was not receptive, the male then would reverse his posture i. e. his anterior now at the posterior tip of the female. After caressing her posterior with his antennae and brushing her anal region with his mouthparts for a few seconds, he would proceed anterior again as he did earlier.

### Mating

After the love play, the male brought his anterior portion of the body to the antero-ventral region of the female while his posterior portion was on her back. Thus the anterior portions of the two bodies were placed ventral to ventral and the male head was enveloping that of the female. In this position ring VII of male lay opposite ring III of the female. Then the male gonopod everted from ring VII inserted into the stretched vulvae of female opened in between rings II/III.

During mating, rapid movement of anterior legs of two or three second duration with intervals of 10, 15, 20, 25 seconds was observed in the male. These rhythmic movement of the legs lasted only for two or three minutes after the start of mating. But no appreciable movement was noticed in the female. At the end of mating or coitus, the male disengaged himself from the female and moved away. But the female in most cases remain stationary in the same place.

In general, when the male came in contact with the female, aggressively overpower the female to mate. It (male) forcefully stretched the anterior portion of the female (often coiled by unwilling female) by holding her head with paired legs of II to V segments. The female however tried to resist such attempts in the very beginning but had to consent ultimately under physical pressure. Such mating was abruptly terminated by the female within two or three minutes.

Sometimes in the field, sexually active solitary males that encounter a mating pair, attempted to take over the female but soon give up and temporarily forms “triplets” for a minute or more.

### Duration of mating

There was considerable variation in the length of the mating period of different pairs in the laboratory as well as in the field. It ranged from one minute which may not have permitted insemination, up to 25 minutes. The average mating took approximately eight minutes.

Two types of mating: long duration and short duration were observed in *C. severini*. The time taken per short duration mating varied from one to six minutes while long duration mating were more than six minutes.

### Mating in relation to time of day

Mating frequency was at its peak in the morning (0600 – 1000) and afternoon (1500 – 1830) (Table 1). Number of mating pairs in the morning is significantly higher than that in the afternoon (P>0.05).

**Table 1.** Number of mating pairs of *C. severini* observed at different hours of the day.

|  |  |  |  |  |  |  |  |
| --- | --- | --- | --- | --- | --- | --- | --- |
| Hour | 06.00 | 07.00 | 08.00 | 09.00 | 10.00 | 11.00 | 12.00 |
| No. of pairs | 89 | 100 | 82 | 66 | 41 | 03 | 02 |
| Percentage | 15.53 | 17.45 | 14.31 | 11.52 | 7.16 | 0.52 | 0.35 |
| Hour | 13.00 | 14.00 | 15.00 | 16.00 | 17.00 | 18.00 | 19.00 |
| No. of pairs | 03 | 02 | 07 | 40 | 95 | 40 | 03 |
| Percentage | 0.52 | 0.35 | 1.22 | 6.98 | 16.58 | 6.98 | 0.52 |

### Effect of month on mating frequency

Mating frequency was maxima in the months of June to September when the millipede were abundant and negligible from December to April (Table 2).

**Table 2.** Number of mating pairs of *C. severini* observed during different months of the year in the field.

|  |  |  |  |  |  |  |
| --- | --- | --- | --- | --- | --- | --- |
| Month | May | June | July | August | September | October |
| No. of mating pairs | 16 | 119 | 123 | 133 | 100 | 46 |
| Percentage | 2.82 | 20.99 | 21.69 | 23.46 | 17.64 | 8.11 |
| Month | November | December | January | February | March | April |
| No. of mating pairs | 16 | 02 | 00 | 00 | 02 | 10 |
| Percentage | 2.82 | 0.35 | 0.00 | 0.00 | 0.35 | 1.76 |

### Mate preference

Result of preference experiment is presented in Table 3. In all cases number of affinities for both sexes is highly significant (P < 0.001).Maximum preference is observed in the female marked as A (Pi = 0.88) while minimum in female C (P_i_ = 0.74) towards the male marked as F and B respectively. Both the male marked as E and F showed maximum (P_i_ = 0.91) and I showed minimum preference (P_i_ = 0.65) towards the female marked as B, A and G respectively. It is observed from the experiment that maximum value of P_i_ depends on i) no mating or very short duration mating ii) maximum number of mating (Table 3). Here preference index for both E and F male is same i. e. 0.91. The male E is mated with only five females out of 10 females but F male though mated with 8 females, but for only one time in all the cases except with female A in which he mated 18 times (Table 3).

**Table 3.** Number of mating (1^st^ row) and duration of mating (in minute, 2^nd^ row) between individually marked *C. severini* for seven days (F = Female, M = Male, N. A. = Number of affinities, P_i_ = Preference index) (X^2^ value, P>0.001).

| F/M | A | B | C | D | E | F | G | H | I | J | N.A. | P<sub>i</sub> | X<sup>2</sup> |
| --- | --- | --- | --- | --- | --- | --- | --- | --- | --- | --- | --- | --- | --- |
| A | 0 | 2<br>3.5 | 0 | 2<br>3.6 | 1<br>2.5 | 18<br>10 | 0 | 0 | 2<br>4 | 0 | 25 | 0.88 | 109.9 |
| B | 1<br>1 | 0 | 4<br>3 | 1<br>4 | 17<br>9 | 1<br>5.5 | 0 | 0 | 1<br>2.4 | 0 | 25 | 0.86 | 98.60 |
| C | 0 | 10<br>8.5 | 0 | 3<br>2.5 | 0 | 1<br>3 | 1<br>3.5 | 4<br>2.5 | 0 | 2<br>3 | 21 | 0.74 | 41.39 |
| D | 1<br>3.5 | 0 | 1<br>3 | 2<br>2.5 | 0 | 1<br>4 | 11<br>9 | 2<br>2 | 3<br>2.5 | 0 | 21 | 0.79 | 46.15 |
| E | 1<br>3.4 | 0 | 2<br>3.2 | 4<br>2.6 | 2<br>3 | 0 | 1<br>2.5 | 9<br>10.4 | 0 | 1<br>2 | 20 | 0.75 | 34.0 |
| F | 1<br>1.5 | 0 | 14<br>9.5 | 0 | 0 | 1<br>1.2 | 1<br>2.5 | 1<br>1.4 | 4<br>5 | 1<br>2 | 23 | 0.82 | 71.33 |
| G | 2<br>2.4 | 1<br>3 | 1<br>2 | 0 | 2<br>2.6 | 1<br>2.4 | 0 | 0 | 9<br>10.2 | 1<br>2 | 17 | 0.83 | 37.71 |
| H | 12<br>8.6 | 0 | 2<br>2.1 | 1<br>1,5 | 1<br>1.5 | 0 | 1<br>2 | 4<br>2 | 0 | 2<br>2.5 | 23 | 0.82 | 51.34 |
| I | 3<br>2.4 | 1<br>2 | 2<br>4.2 | 1<br>2 | 0 | 1<br>1.5 | 0 | 1<br>1 | 1<br>2 | 8<br>10.4 | 18 | 0.78 | 27.58 |
| J | 0 | 1<br>3 | 0 | 10<br>9.4 | 0 | 1<br>2.2 | 0 | 1<br>2 | 4<br>2.5 | 2<br>2 | 19 | 0.82 | 45.75 |
| N.A. | 21 | 15 | 26 | 24 | 23 | 25 | 15 | 22 | 24 | 17 |  |  |  |
| P<sub>i</sub> | 0.83 | 0.85 | 0.79 | 0.71 | 0.91 | 0.91 | 0.90 | 0.78 | 0.65 | 0.80 |  |  |  |
| X<sup>2</sup> | 55.68 | 56.35 | 60.91 | 32.69 | 106.99 | 107.40 | 68.35 | 32.53 | 29.35 | 29.47 |  |  |  |

## Discussion

*C. severini* follows the basic pattern of love play described for other Polydesmids (Seifert, 1932; Causey, 1943; Mauries, 1969; Snider, 1981, Mukhopadhyaya and Saha, 1981). In all the cases, male approaches the female along her dorsum, draws the female around so that ventral surfaces of the pair become contagious and the male held the female’s anterior portion tightly by paired legs.

Triplet formation by sexually active solitary male is also observed in millipede, *Centrobolus inscriptus* (Cooper and Telford, 2000) and in *Odontota dorsalis* (Kirkendall, 1984). Telford and Dangerfield (1993a) suggested in a highly male biased situation, triplet formation can occur if a male came in contact with a copula randomly. In the present study, triplet formation in *C. severini* follows the same rule.

In millipede, duration of mating has been reported to last from 10 minute to over 48 hours (Schubart, 1934; Stephenson, 1961; Sahli, 1964; Mauries, 1969; Snider, 1981; Mukhopadhyaya and Saha, 1981; Bhakat et al.1989; Holwell et al., 2016). Generally in Polydesmida, mating duration is comparatively low viz. 1 to 15 min. (*Orthomorpha coarctata*), 2.6 to 17.2 min. (average 4.7 ± 2.7 min.) (*Cladethosoma clarum*), 10.38 ± 0.86 min.(*Gigantowales chisholmi*), 14.5 min. (*Streptogonopus phipsoni*) but in other orders mating duration is too long viz. >4 hours (*A. uncinatus*), >2 hours (*Calostreptus* sp.), >2 hours (*Spinotarsus* sp.) >3 hours (*Centrobolus inscriptus*). Being a Polydesmid, *C. severini* follows the general rule of short mating time which ranges from 1 min. to 25 min. with an average of 8 min.

Mukhopadhyaya and saha (1981) observed two types of mating in *O. coarctata* – short and long duration. Time taken for long duration mating is 15 min. preceeded by a short love play or without it, while that of short duration mating ranges from 1 to 8 min. preceeded by a longer love play. In the later case, female shortened the time by forcibly disengaged herself from the male. They also linked short duration mating with repeated copulation. In the present study, both in the field and in the multiple choice experiment, two types of mating was also observed – short duration (maximum 6 min.) and long duration (maximum 25 min.). But in both types, duration of love play does not varied significantly. The duration of mating is a measure of the period of genitalic contact between mating partners (Sillen and Tullberg, 1981). The short duration mating were recorded for species that showed less vigorous mating as observed in Juliform millipedes (Telford and Dangerfield, 1990; Cooper and Telford, 2000) where differences in mating time are thought to consider the intensity of sperm competition between species. Holwell et al. (2016) argued that short duration copulation reflect a general mating strategy that emphasizes mate acquisition over mate guarding. In diplopods, long duration mating is presumably an adaptation in the form of mate guarding and sperm competition (Heisler, 1983; Telford and Dangerfield 1990, 1993a, b; Barnett and Telford, 1996; Rowe, 2010; Cooper, 2016). But till today, sperm competition or mate guarding in millipede is not quantified rather predictions are made by passive observation of mating behaviour of males and females. Moreover, a few authors (Eady, 1994; Kelly and Jennious, 2011) commented that copulation duration does not necessarily correlate with sperm transfer or fertilization success. Some also argued that a male’s investment in copulation duration may be traded off against his likelihood of achieving future mating success (Andrade, 2003; Kasnmovic et al. 2007; Rowe, 2010). In this context Simmons’s (1991) comment is quite relevant here. He suggested that mean copulation duration for a species at any one instant should be considered as an outcome of sexual conflict within pairs which depends on the relative ability of both sexes to exert their interest over one another. In *C. severini* though short and long duration mating is initiated by male but their duration may depend on the conflict of participating male and female which is presumably controlled by several factors like gonopod structure, size of mate, pre-copulatory time gap, maturity of female or other stimuli. I also think that long duration mating is an adaptation in the form of mate guarding and also an advantage for the female in the form of maximum fertilization of ova which increase their reproductive success and/or fitness while short duration mating is a process of mate acquisition in a competitive way.

Like other millipedes, mating frequency of *C. severini* is maximum in the morning (58.81%) and afternoon hours (30.54%) and in the months of fall i.e. June to October (91.89%) (Demange, 1959; Banerjee, 1973; Rangaswami, 1973; Mukhopadhyaya and Saha, 1981; Mukhopadhyaya and Bhakat, 1983).

During breeding season, multiply mating in millipede is a very common phenomenon in millipedes (Mukhopadhyaya and Saha, 1981; Carey and Bull, 1986; Tadler, 1993; Telford and Dangerfield, 1993a; Rowe, 2010; Wojcieszek and Simmons, 2011; Holwell et al. 2016). Multiple or repeated mating has been observed in *O. coarctata* where the species copulated six times in 24 hours and duration of each mating lasted from 10 to 12 min. Holwell et al. (2016) reported multiple mating in a Polydesmid, *Gigantowales chisholmi*. In this species male follows a scramble competition strategy. The male repeated mate with females for a brief period of time without further attempt to a particular female. In this way a single male can mate with numerous female within a short period of time. In the present study, in multiple choice experiments, both male and female *C. severini* mated multiple times with short and long duration mating. It seems that both sexes can exert pre-copulatory as well as cryptic choice and previous experience may influence mate choice in this species. Like various arthropods (Zeh et al. 1998; Li et al. 2014), preferences for previous or new mates in choice test suggest a possible linkage with their different mating demands and strategies. Like other Polydesmids (Blower, 1970; Bhakat, 2014), *C. severini* is semelparous i.e. lays eggs for one time. So the female needs more sperm to fertilize maximum number of ova which is only possible through multiple mating or very long duration mating as found in Julida, Spirobolida and Spirostreptida (Carey and Bull, 1986; Tadler, 1993; Telford and Dangerfield, 1993a; Barnett, 1997). As the duration of mating in *C. severini* is not very long (compared to other non-polydesmid millipede), so the only option for maximum fertilization is multiple mating. The process of multiple mating is a form of sexual competition i. e. the competition between males for access to females or competition between males to be chosen by females (Darwin, 1871; Patridge and Halliday, 1984). Potential benefit to females from multiple mating is that it increases female survival, fecundity and fertility (Yasui, 1998; Arnqvist and Nilsson, 2000; Hosken and Stockley, 2003) as well as increases the genetic diversity of offspring (Yasui, 1998; Slatyer et al., 2012).

In conclusion, the study confirms that *C. severini* is characterized by a polygynandrous mating system where males are the pursuers and females are the accomplisher of mating and duration of mating for short and long period related to mate acquisition and mate guarding respectively.

## Acknowledgement

I am indebted to my son Dr. Soumendranath Bhakat, Lund University, Sweden for his help and grateful to all my colleagues for their inspiration.

